# Root shape phenotyping using three-dimensional root system vector data in rice

**DOI:** 10.64898/2026.06.15.732232

**Authors:** Shota Teramoto, Yusaku Uga

## Abstract

**Purpose:** Alternation of root distribution in the soil is a method for improving root system architecture (RSA) in crops. If the root is straight, root distribution has been altered by regulating the angle of root growth in the vertical direction. However, the root shape should be curved and winding. This study aimed to define parameters reflecting the actual root shape.

**Methods:** We used three-dimensional vector data of rice *(Oryza sativa* L.) RSA derived from an X-ray computed tomography image to compile two sets of two-dimensional vector data for horizontal and vertical components. In the vertical component, we defined the dropping angle *θ*_*d*_, which is calculated assuming that the rice roots are bent upward. In the horizontal component, we defined the polar angle *θ*_*ρ*_, which is the direction in which the roots grow when viewing the plant from above, and the winding degree log σ_*w*_, which is calculated by assuming that root elongation is in a random walk.

**Results:** Assuming that *θ*_*ρ*_ is distributed uniformly in all varieties, there should be no varietal differences in this angle. We measured *θ*_*d*_ and log σ_*w*_ of three rice varieties with different root distributions: shallow, deep, and intermediate. We found significant varietal difference in *θ*_*d*_ and log σ_*w*_.

**Conclusions:** We have shown that *θ*_*d*_ and log σ_*w*_ are useful parameters for comparing RSA in rice varieties. We have named this methodology RSAparam3D, and it is freely available to researchers.

## Background

Root system architecture (RSA) refers to the spatial distribution of various types of roots in the soil (Lynch 1995). In monocots, RSA is the fibrous root system comprising a radicle, crown roots, and lateral roots (Steffens and Rasmussen 2016). For example, in rice *(Oryza sativa* L.), the radicle is the embryonic root that first emerges from the seed (Itoh et al. 2005); it has many lateral roots and plays important roles in the plant taking root and growing during the juvenile stage. Crown roots are nodal roots that have many lateral roots and constitute the main skeleton of a fibrous root system (Itoh et al. 2005). RSA determines the availability of nutrients and water distributed heterogeneously in the soil (Lynch 1995; de Dorlodot et al. 2007). Therefore, RSA is an important crop characteristic that greatly influences productivity (de Dorlodot et al. 2007; Thorup-Kristensen et al. 2020). RSA modification is considered to be a useful approach to facilitate crop improvement (Uga et al. 2015a; Uga 2021). However, carrying out RSA improvement is difficult and laborious because it requires considerable effort to dig up roots from the soil (Böhm 1979). Moreover, digging up roots from the soil often causes loss of three-dimensional (3D) information such as root distribution in the soil (Trachsel et al. 2011).

The RSA skeleton is determined by four main parameters: growth, branching, surface area, and angle (Morris et al. 2017). In monocots, these four parameters are determined by root number, root growth angle (RGA), and root density. Root number determines growth and surface area, RGA determines angle, and root density determines growth, branching, and surface area. Generally, root number indicates the number of nodal roots (Obara et al. 2014; Zhang et al. 2018), RGA indicates an angle of vertical nodal root elongation when the horizontal is 0° (Oyanagi et al. 1993), and root density indicates total root length including lateral roots per unit area (Yoshida and Hasegawa 1982). Of these, RGA alternation is employed widely for RSA improvement (Uga et al. 2013; Kitomi et al. 2020) because RGA exerts a considerable effect on root distribution in the soil (Abe and Morita 1994) and is easy to measure with appropriate equipment (Oyanagi et al. 1993), such as a mesh basket. This procedure entails, planting the plants in soil-filled baskets and measuring RGA by connecting the planting point and the point where the roots protrude from the mesh with a straight line (Oyanagi et al. 1993; Kato et al. 2006). However, the actual shape of the roots should be curved and winding, not straight (Morris et al. 2017). There are few examples of such architecture being measured because simple methods for measuring 3D shapes are lacking. Accurate quantification of the root shape enables a more detailed evaluation of various characteristics.

In recent years, information on the 3D deployment of roots in the soil has become easy to obtain using X-ray computed tomography (CT), magnetic resonance imaging, and other techniques (Atkinson et al. 2019). For phenotyping, root density (van Dusschoten et al. 2016) or root length (Gao et al. 2019b, a) in each soil layer is generally calculated by isolating root segments from 3D images because root segmentation is relatively easy to carry out. However, segmentation cannot represent the root shape. Another phenotyping method with a 3D volume image is root vectorization, which represents root shape as a vector; any position on a root has corresponding sets of coordinates (Lobet et al. 2011; Teramoto et al. 2021). We assumed that we could use 3D vector data to measure not only RGA calculated based on the roots being straight (Teramoto et al. 2021) but also root shape, whether straight, curved, or winding. This study demonstrates comparison of the differences in RSA by extracting parameters that represent 3D shapes from 3D vector data of rice RSA.

## Materials and methods

### Plant materials

In this study, we used three rice varieties: IR64 (IRGC #66970), Kinandang Patong (KP, IRGC #23364), and Dro 1 -NIL (Uga et al. 2013; Teramoto et al. 2019). KP has deep RSA, whereas IR64 has shallow RSA. Dro1 -NIL is a near-isogenic line of IR64 with a KP-type *DRO1* allele, and it has intermediate RSA.

### Growth conditions

We adopted the growing conditions used by Teramoto et al. (2020). We sowed rice plants in pots measuring 25-cm depth and 20-cm diameter (TSP2530P, Tecs, Itako, Ibaraki, Japan) filled with a soil-like root growth substrate (Profile® Greens Grade^™^, PROFILE Products, Buffalo, Illinois, USA) and cultivated the plants in a growth chamber for 4 weeks. Before sowing, we saturated the substrate with a hydroponic solution and supplied deionized water from the bottom of the pots during cultivation. The hydroponic solution consisted of 1.23 mM of NO_3_^-^, 0.41 mM of NH_4_^+^, 0.18 mM of H_2_PO_4_^-^, 1.00 mM of SO_4_^2-^, 1.78 mM of K^+^, 0.55 mM of Mg^2+^, 0.37 mM of Ca^2+^, and 8.9 μM of Fe^3+^. We adjusted the pH to 5.5, using HCl and KOH, as required. The light conditions we used simulated 14 h of daylight. Details of other conditions have been described in Teramoto et al. (2020). To buffer the positional effect, we adopted a block design. Six blocks were used, with each block having three pots: IR64, Dro1-NIL, and KP.

### X-ray computed tomography scanning

We adopted the X-ray CT scanning and root segmentation conditions used in Teramoto et al. (2020). We imaged rice roots in the soil using the X-ray CT system inspeXio SMX-225CT FPD HR (Shimadzu Corporation, Nakagyo-ku, Kyoto, Japan). Each scan was obtained digitally; 1,200 projections, using a signal averaging two frames over 360° without binning (pixel detector resolution: 3,000 × 3,000), at 4.0 fps. We computed 860 horizontal, 1,024 × 1,024 pixel-resolution slices. The final spatial resolution was 300 pm, corresponding to a total volume of 30.72 × 30.72 × 25.8 cm^3^. The tube voltage was 225 kV, and the tube current was 500 μA. Source-detector and -rotation axis distances were 1,200 and 900 mm, respectively. To harden the X-ray beam, we used a 1 .0-mm Cu filter. Beam hardening was corrected approximately using the operating software, with a correction table calculated using the correct metal material.

### Root segmentation and vectorization

We imaged root segments in X-ray CT images using RSAvis3D (Teramoto et al. 2020). The parameters of RSAvis3D were as follows: median filter size was 7, edge detection filter size was 21, and cylinder radius was 9 cm. We vectorized segmented volume data using the “COG tracking” method in RSAtrace3D (Teramoto et al. 2021). We saved vectorized data in a JavaScript Object Notation (JSON) format.

### Parameter estimation for vertical root system architecture phenotype

Given that the i-th coordinate of a vector is *c*_*i*_-(*x*_*i*_-,*y*_*i*_-*z*_*j*_), where *x*_*i*_. *y*_*i*_. *z*_*i*_ are Euclidean coordinates for the *x. y*. and *z* axes. respectively. and that *n* is the number of coordinates in the vector. the vector is Eq(1).

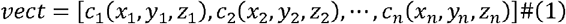

Based on Eq. (1). the i-th vertical local angle *θ*_*vi*_ is defined as Eq. (2).

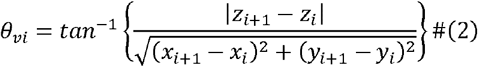

Given that the dropping angle is *θ*_*d*_ and the decay rate is *a* (0.005). the modeled local angle *θ*_*vsi*_ is represented as Eq. (3).

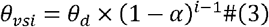

Based on Eqs. (2) and (3). *θ*_*d*_ is determined to minimize Eq. (4) using Python3 (https://www.python.org/) with the “least_sequence” function in the “scipy.optimize” module (Virtanen et al. 2020).

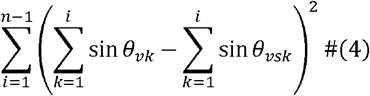

### Parameter estimation for horizontal root system architecture phenotype

Based on Eq. (1). the i-th horizontal local angle *θ*_*hi*_ (0^°^ < *θ*_*hi*_ < 360°) is defined as Eqs. (5) and (6).

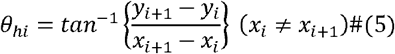

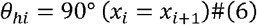

Based on Eqs. (5) and (6). Δ*θ*_*hi*_; (|*θ*_*hi*_| *I* < 180°) is defined as Eq. (7).

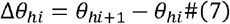

For Δ*θ*_*hi*_, where *i* is from 1 to *n-1*. we calculated winding degree *σ*_*w*_ using Python3 with the “norm.fit” function in the “scipy.stats” module (Virtanen et al. 2020).

### Statistical analysis

We carried out Student’s t-tests and Tukey-Kramer tests using R software (https://www.r-project.org/) and the “multcomp” package (Bretz et al. 2016). A *P* valued<0.05 was considered significant.

## Results

The overall diagram of the phenotyping method with vectorized RSA data is shown in Fig. 1. Vectorized 3D RSA comprised the vectorized radicle and crown roots (Fig. 1a). We defined the direction of root elongation as the longitudinal axis and the axes horizontally and vertically perpendicular to the longitudinal axis as the horizontal and vertical axes, respectively (Fig. 1b). To simplify the model, we decomposed the 3D data into two sets of 2D data (Figs. 1c and d). To define the vertical movement of roots, we offset the movement along the horizontal axis (Fig. 1c). One end of each root was gathered at the center of the image (Fig. 1c). We constructed a vertical 2D root model using two parameters: the direction of root growth from the center (similar to the concept of RGA) and bending degree of the roots. To define the horizontal movement of roots, we offset the movement along the vertical axis (Fig. 1d). The root is represented by a single curve connecting the center and end of the root (Fig. 1d). We constructed a horizontal 2D root model using two parameters: the direction of root growth from the center (polar angle) and winding degree of the left-right wobble of the root shape. Finally, we calculated frequency distributions representing the RSA phenotype (Figs. 1c and d).

**Fig. 1.**
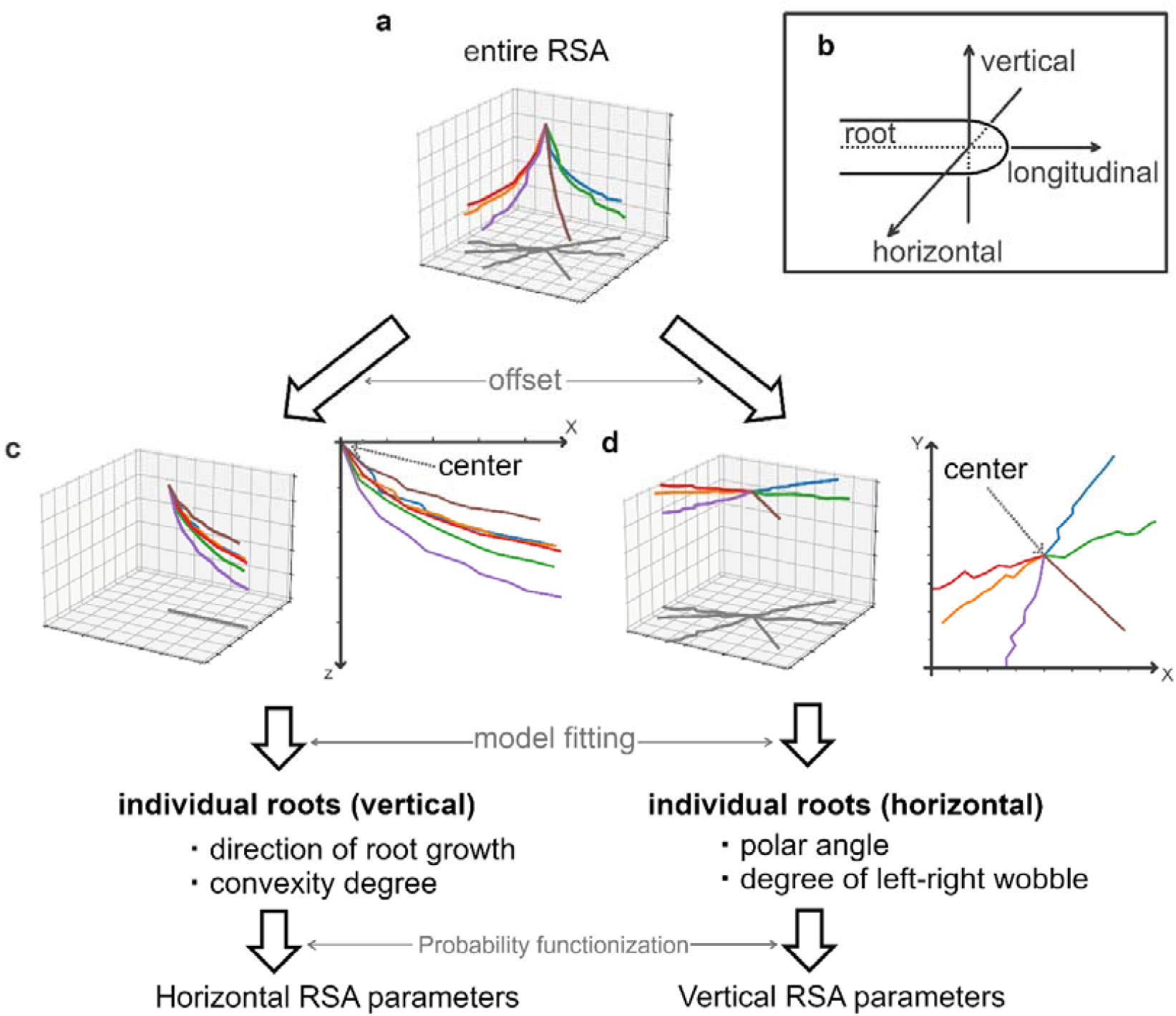
Root system architecture (RSA) phenotyping methodology from vectorized rice RSA. **a** Three-dimensional rendering of vectorized rice RSA. Each root is colored. **b** Definition of the axis for root movement. Three-dimensional rendering of vectorized rice RSA (left) and a two-dimensional line graph (right): **c** horizontal movement offset; and **d** vertical movement offset

### Parameters for vertical root shape

We used vectorized RSA data of an upland rice variety, KP (Teramoto et al. 2020, 2021). Each root is represented by an unbranched vector with the distance between nodes being approximately four voxels. To parameterize the vertical root shape, we converted the 3D RSA vector to a 2D vector by offsetting the horizontal axis (Fig. 2a). Most roots extended downward to the right, with a slight upward bend (Fig. 2a). We defined an angle against the ground surface of each node as the local angle and assumed the root shape to be represented by the initial local angle and decay rate *a*, whereby the local angle decays by *a* over nodes to a small value close to zero (Fig. 2a). We converted the vector data to node index-depth data (Fig. 2b left) and then estimated the initial local angles by fitting with mean square error as a loss function, with 0.005 of *a* (Fig. 2b right). We defined the fitted initial local angle as a vertical root shape parameter, the dropping angle *θ*_*d*_.

**Fig. 2.**
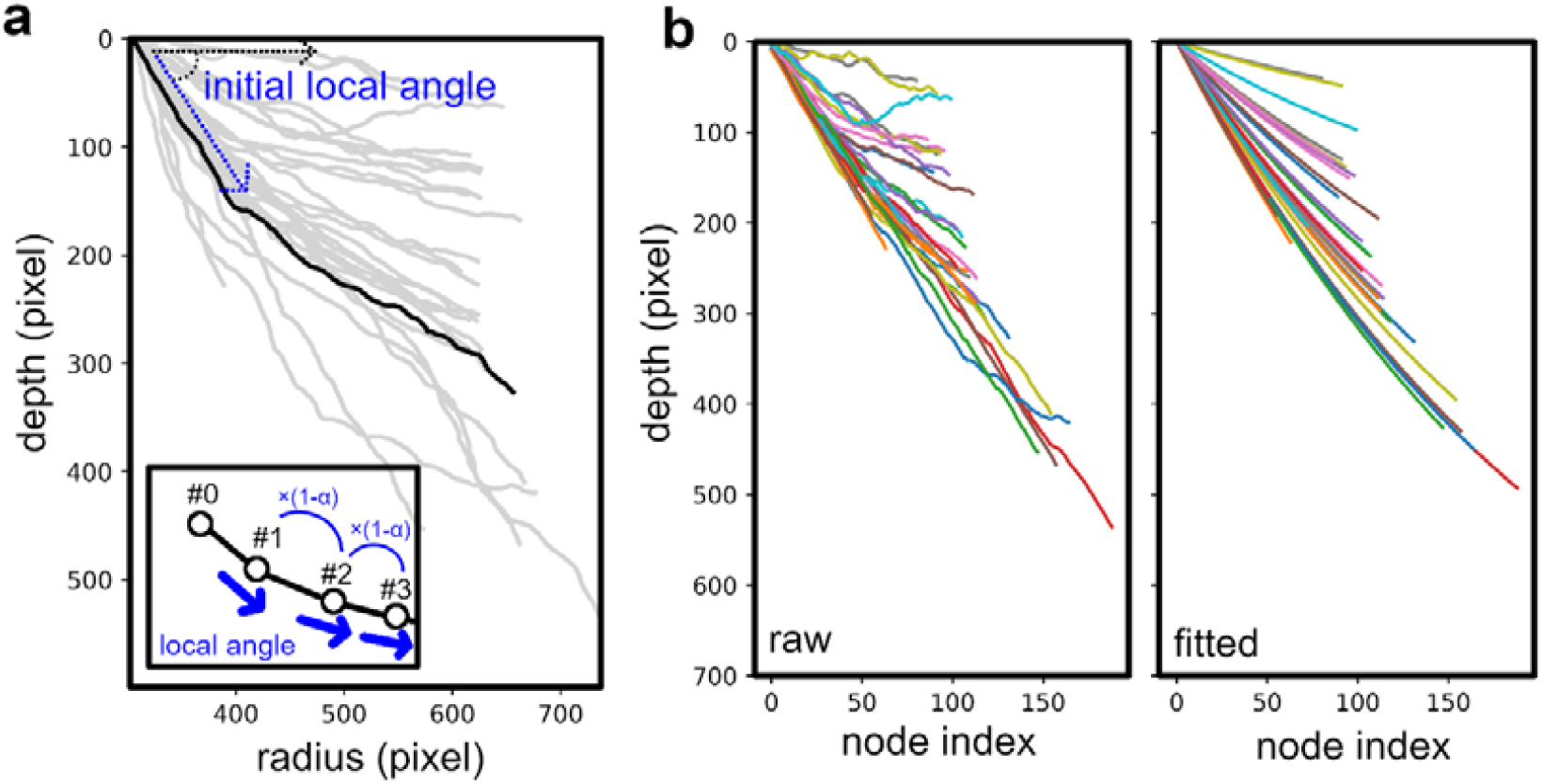
Fitting of vertical movement of vectorized rice root system architecture. **a** Line graph with horizontally offset data. **b** Line graph with data converted into node index and depth (left), and line graph with data generated by the fitted model (right)

### Parameters for horizontal root shape

To parameterize the horizontal root shape, we converted the 3D RSA vector to a 2D vector by offsetting the vertical axis (Fig. 3a). We defined an angle on the x-y plane of each node as the local angle (Fig. 3a) and the initial local angle as the polar angle *θ*_*p*_. We calculated the normalized local angle by offsetting the *θ*_*p*_ (Fig. 2b) from the local angles of all nodes. The curve in Fig. 3b took the form of a random walk, likely because the roots elongate by random bumps into soil particles. As the random walk is represented by the accumulation of random numbers, we assumed that the standard deviation of a random number should be a parameter representing the horizontal root shape. We calculated the change in the normalized local angle and its frequency distribution (Figs. 3c and d). We obtained a parameter *σ*_*w*_ as standard deviation of the normal distribution (Fig. 3d). The parameters obtained—*θ*_*p*_ and *σ*_*w*_*—*represent the horizontal root shape.

**Fig. 3.**
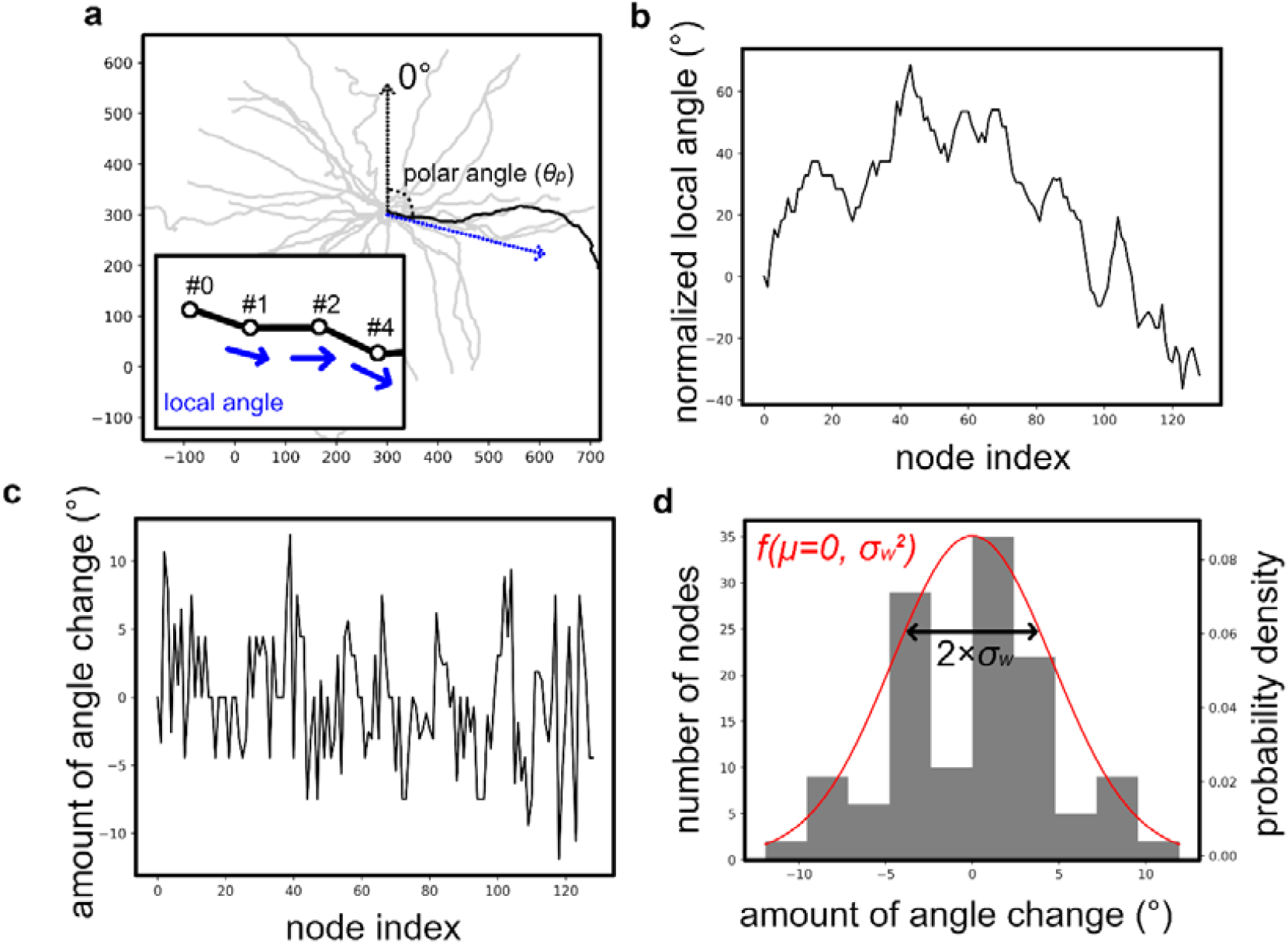
Fitting of horizontal movement of vectorized rice root system architecture. **a** Line graph with vertically offset data. **b** Line graph with data converted into a node index and local angles of nodes normalized so that the polar angle is zero. **c** Line graph with node index and amounts of normalized local angle changes. **d** Histogram with amounts of normalized local angle changes fitted to a normal distribution with mean 0 and variance *σ*_*w*_

### Root system architecture phenotyping based on the frequency distribution of root parameters

To ascertain whether the obtained parameters can be used to phenotype root shape, we parameterized three RSA types: IR64, Dro1-NIL, and KP. In IR64, the roots are concentrated close to the surface of the ground, producing a shallow-rooting RSA phenotype, whereas KP has a deep-rooting RSA phenotype (Uga et al. 2011). Dro1-NIL is a near-isogenic line of IR64 with a KP-type allele of the *DEEPER ROOTING 1 (DRO1)* gene that controls the angle of root growth (Uga et al. 2013). Dro1-NIL has intermediate RSA types of IR64 and KP (Uga et al. 2015b; Teramoto et al. 2019). IR64, Dro1-NIL, and KP are suitable for testing the phenotyping capacity of the model in this study because previous studies have used them as three different representative varieties of RSA (Teramoto et al. 2019; Teramoto and Uga 2020). The vector data we use in this study are illustrated in Fig. 4.

**Fig. 4.**
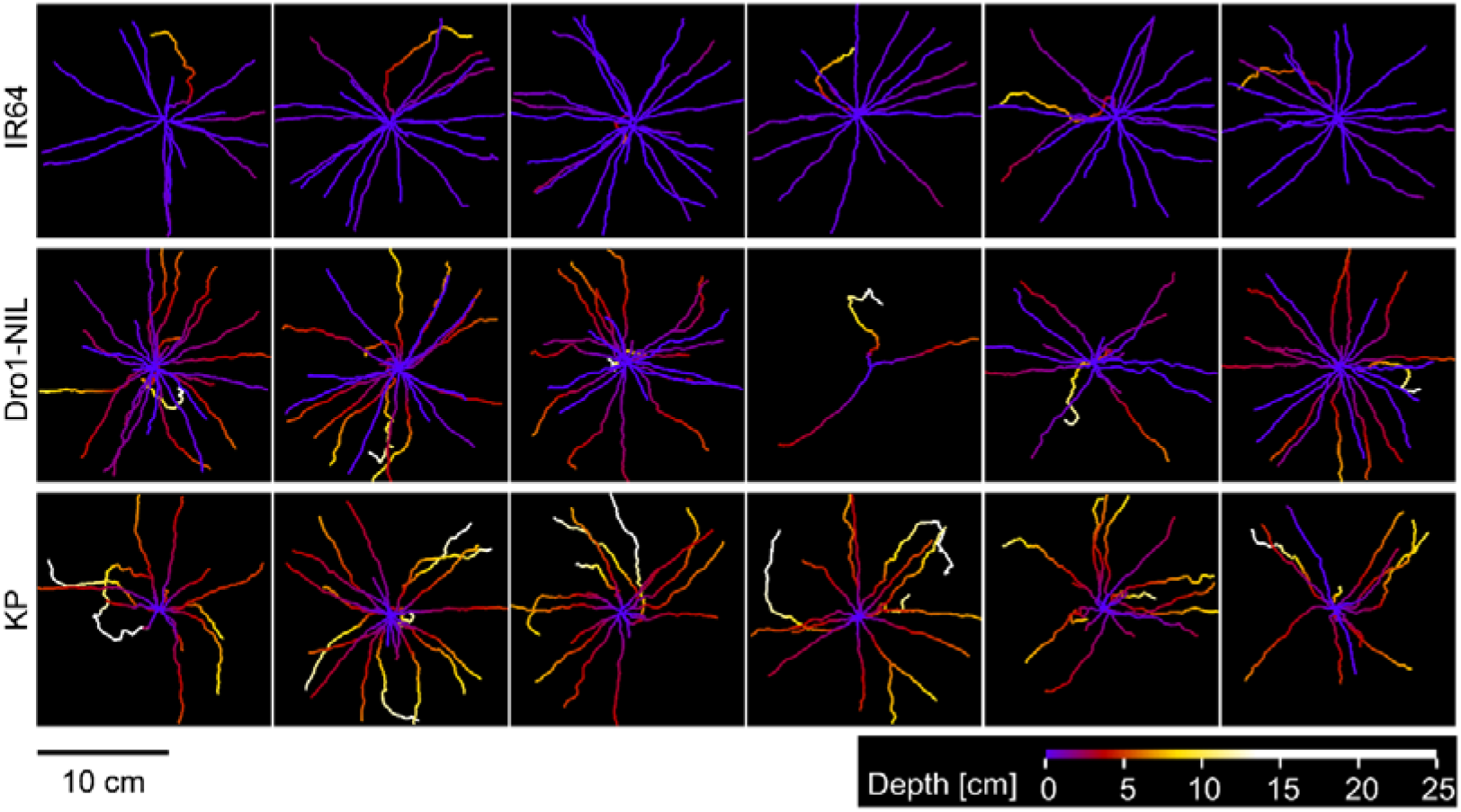
Vector data of root system architecture in IR64, Dro1-NIL, and Kinandang Patong. The overhead view is illustrated. The color scale represents the root depth when the depth at which the seed was planted is set to zero

In terms of the dropping angles, *θ*_*d*_, the histogram of KP appears symmetrical, whereas those of IR64 and Dro1-NIL do not appear symmetrical because of the large numbers of *θ*_*d*_ of approximately 0° (Figs. 5a-c). As the roots of IR64 and Dro1-NIL are close to the surface of the ground, they are likely to have more *θ*_*d*_ of approximately 0°. In each of the three RSA types, the *θ*_*d*_ of the radicle was apparently higher than was the *θ*_*d*_ of the crown roots. In terms of winding degrees, *σ*_*w*_, the *σ*_*w*_ of the radicle was also apparently higher than was the *σ*_*w*_ of the crown roots in the three RSA types (Figs. 5d-f). The distributions were long-tailed in shape (Figs. 5d-f), indicating that they were close to log-normal distributions. We log-transformed the *σ*_*w*_ and adjusted the long tails to be close to symmetrical (Figs. 5g-i).

**Fig. 5.**
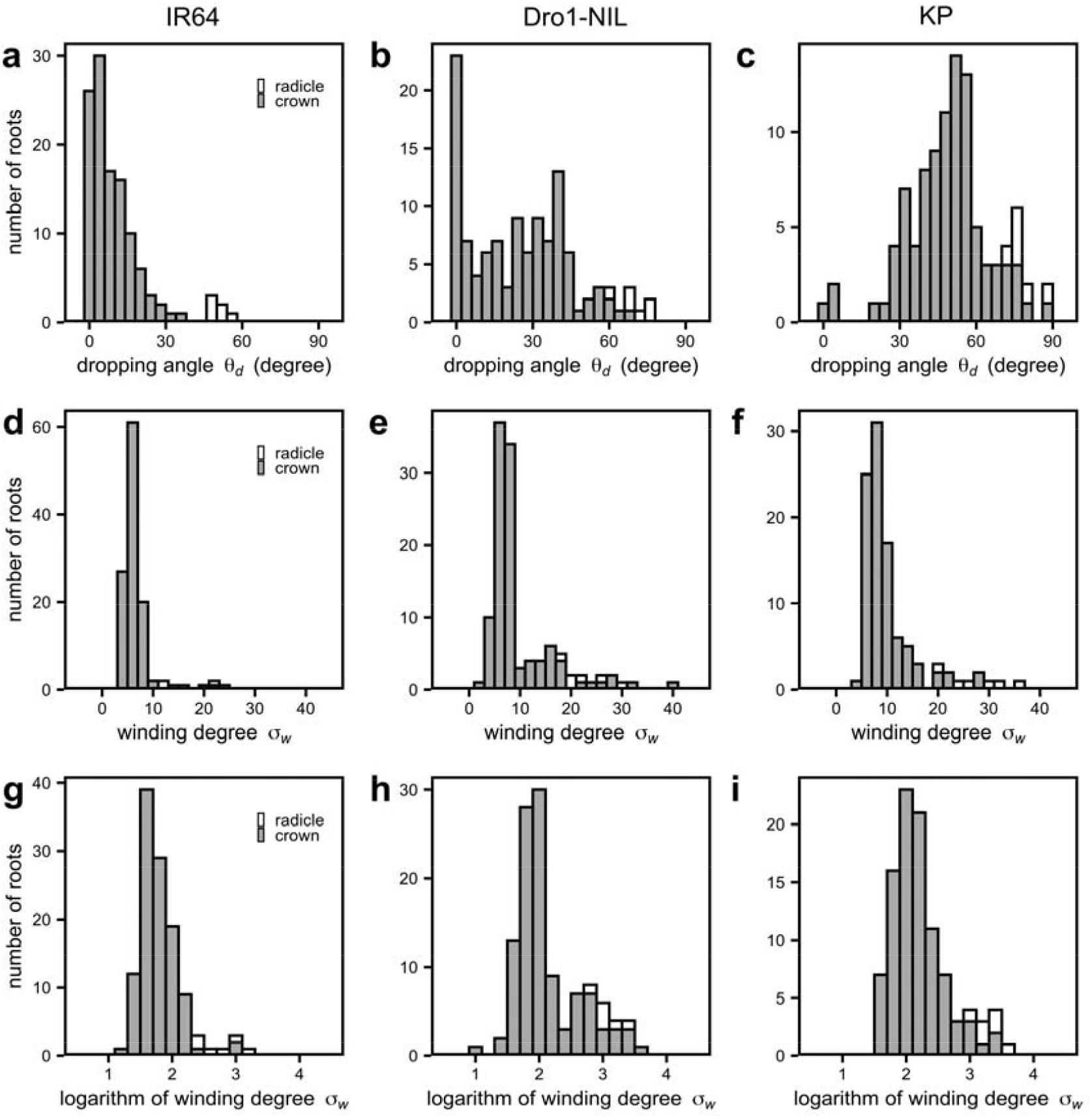
Frequency distribution of root parameters. Six individuals of each variety were bulked. Histograms of radicle and crown roots: **a-c** dropping angles *θ*_*d*_. **d-f** winding degrees σ_*w*_; **g-h** logarithmic winding degrees log σ_*w*_; **a, d, g** IR64; **b, e, h** Drol-NIL; **c, f, i** Kinandang Patong. Six individuals of each variety were used

We calculated the horizontal and vertical RSA parameters—mean of 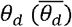 and mean of 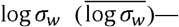of IR64, Dro1-NIL, and KP (Fig. 6). The 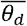 and 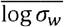 of the radicles were significantly higher than those of the crown roots (Figs. 6a and b), indicating that the radicle elongates more deeply and has greater horizontal movement. In both vertical (Fig. 6a) and horizontal parameters (Fig. 6b), 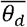 and 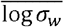 of IR64 in both crown roots and radicles were significantly lower than those of KP (Fig. 6a, b). Dro1-NIL was the intermediate type and 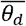 and 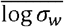of IR64 were significantly lower than those of Dro1-NIL in both radicles and crown roots (Figs. 6a and b), meaning that the *DRO1* gene may not only affect the angle of growth of the vertical root but also horizontal movement. However, we found a correlation between 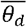 and 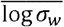, suggesting that the deeper the root, the greater the horizontal movement (Fig. 6c). Further experiments are required to ascertain the relationship between 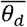 and 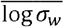. Using RSA parameters of each variety, we simulated the RSA of each (Fig. 7). The simulation yielded shallow, deep, and intermediate RSAs, indicating that the measured parameters reflect the varietal characteristics.

**Fig. 6.**
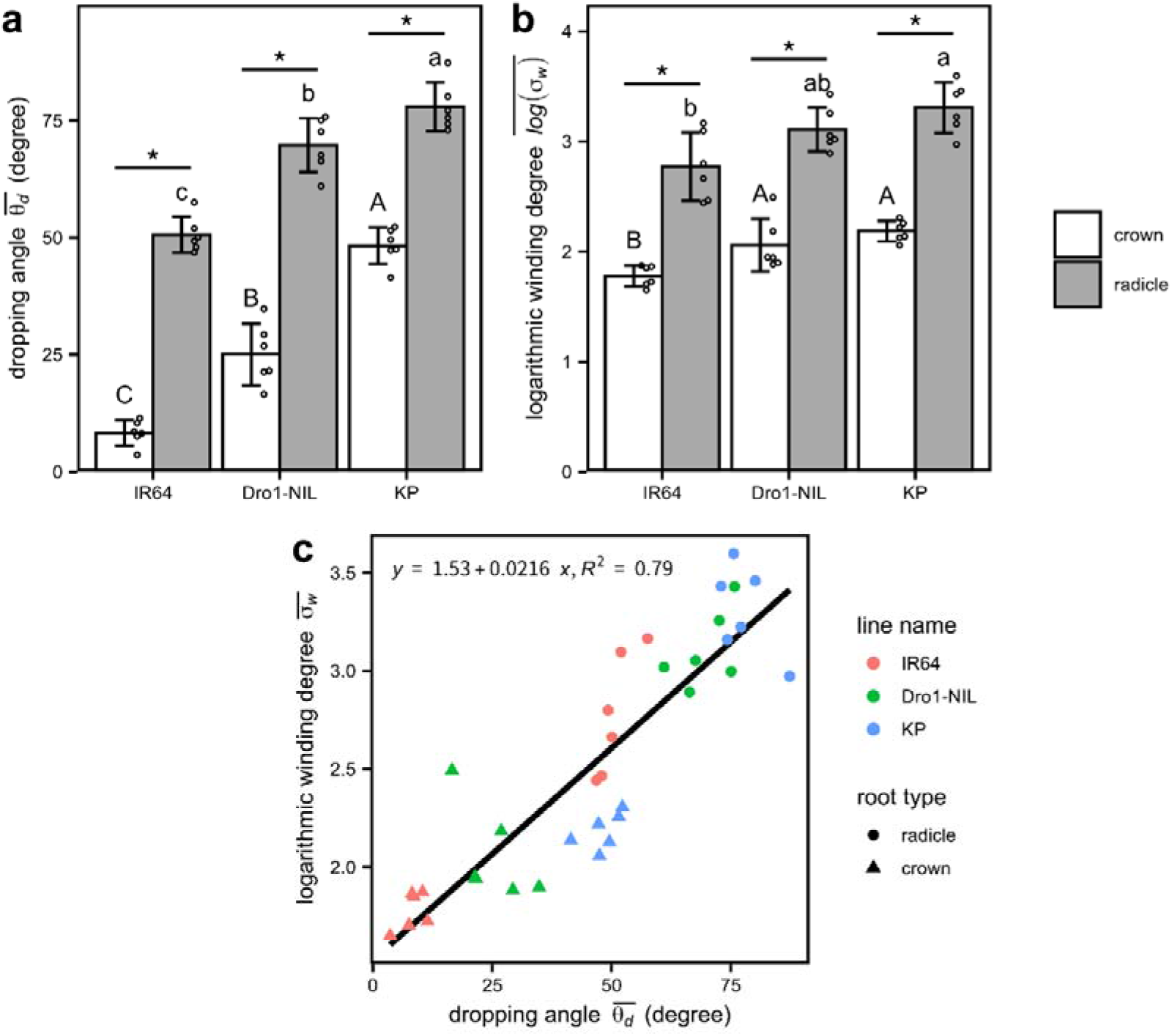
Comparison of 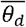and 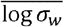 of IR64, Dro1-NIL, and Kinandang Patong. Six individuals of each variety were used. Bar graphs of radicles and crown roots; **a** mean of 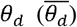: **b** mean of log σ_*w*_ 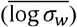. The letters above the bars indicate a significant difference calculated using the Tukey-Kramer test. *: significant difference (Student’s t-test). The error bars indicate the standard deviation. The raw data making up the bar graph are indicated by small circles. **c** A scatter plot of 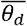 and 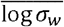. The straight line was drawn using a linear regression

**Fig. 7.**
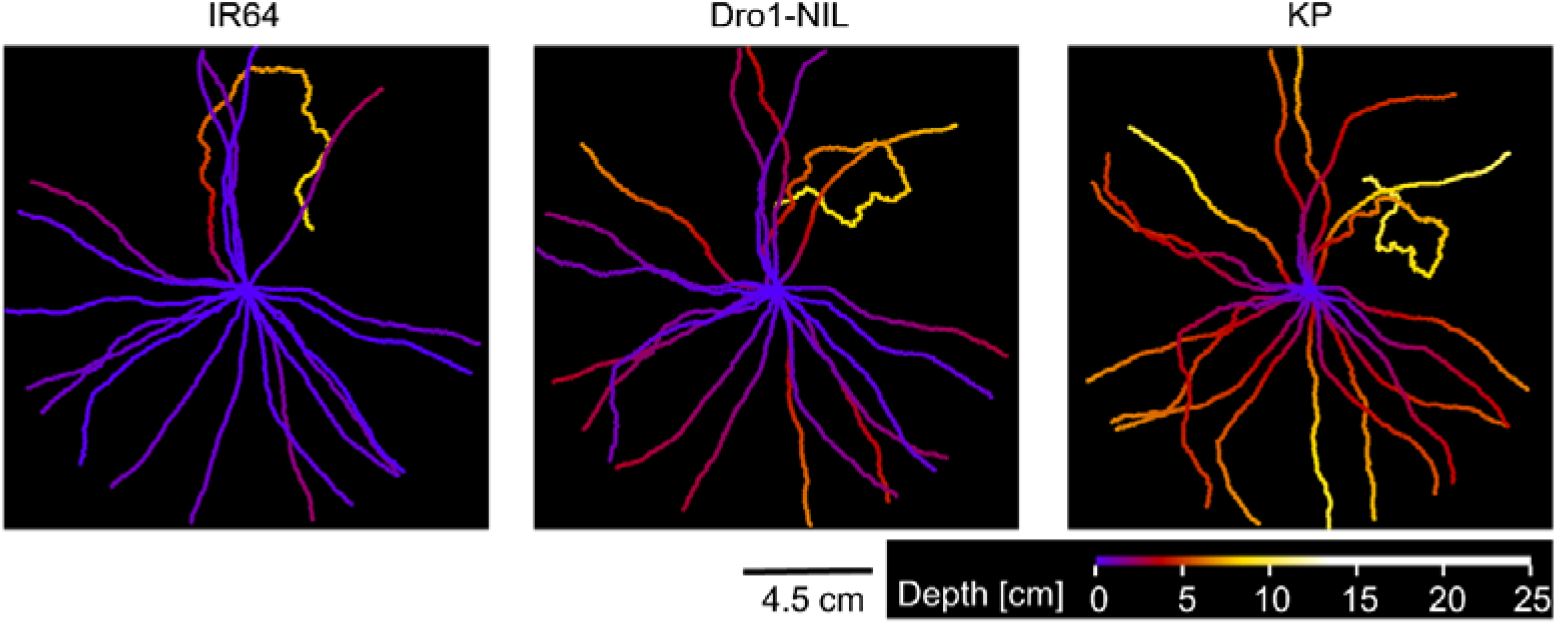
Simulated root system architecture with the measured parameters. Twenty-five roots were simulated

### Root growth angle is influenced by the horizontal parameter

RGA is affected by a range of factors including soil compaction (Correa et al. 2019), temperature (Nagel et al. 2009), and root diameter (Materechera et al. 1992). In this study, we examined the effect of the horizontal parameter log σ_*w*_ and distance in RGA measurement (distance between the planting site and the point of measurement of RGA [Fig. 8a]), which is considered to have an impact, combined with log σ_*w*_, on RGA. Fig. 8b shows the effect of distance in RGA measurement and log σ_*w*_ on RGA estimated via simulation, and the raw data are provided in Online Resource 1. We conducted the simulation with *θ*_*d*_ fixed at 60°. Irrespective of log σ_*w*_, when distance in RGA measurement is 0, the RGA was simulated as 60°. As the distance in RGA measurement increased, the RGA was simulated smaller, reducing RGA to 40°. This is because the roots are bent upward (Fig. 2). Conversely, when log σ_*w*_ was high, the RGA was large. In Fig. 8 the maximum RGA is 69°. This was because undulation of the roots increased the time taken to reach the length to measure RGA. To summarize, the calculation of RGA assuming the roots are straight is affected by both distance in RGA measurement and log σ_*w*_, and *θ*_*d*_, which eliminates both effects, is a more accurate vertical parameter.

**Fig. 8.**
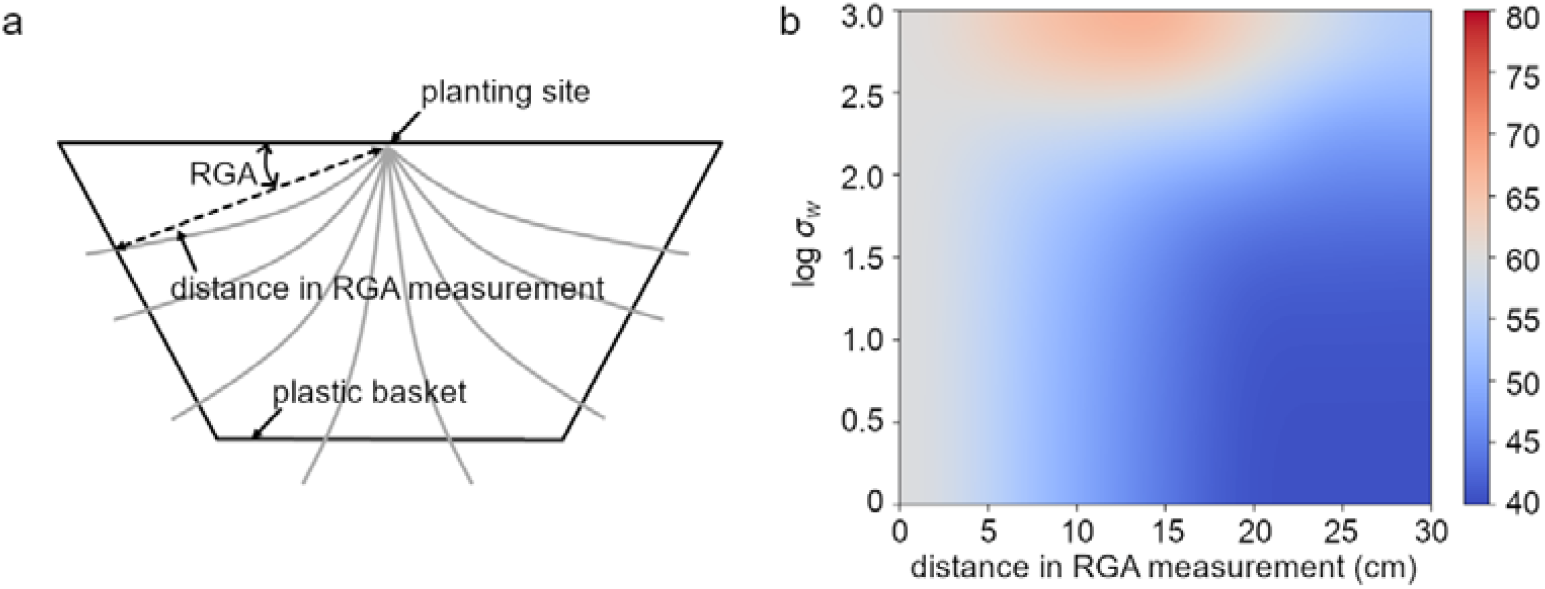
Influence of distance in RGA measurement and 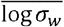. **a** Schematic diagram of RGA measurement. When measuring the RGA, the roots are assumed to be straight. **b** Simulation analysis of the effect of distance in RGA measurement and 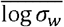 on RGA. *θ*_*d*_ was fixed at 60 degrees. The plots were smoothed by the “bicubic” method

## Discussion

Three-dimensional RSA phenotyping is one of the key processes in modification of RSA in crops. In this study, we quantified the 3D vectorized RSA of rice by using three parameters—dropping angle *θ*_*d*_, polar angle *θ*_*ρ*_, and logarithmic winding degree log σ_*w*_—to describe the shapes of the radicles and crown roots. Assuming that *θ*_*ρ*_ is distributed uniformly in all varieties, the varietal differences in RSA are explained by *θ*_*d*_ and log σ_*w*_. We defined the means of 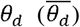 and 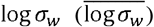 in each individual as representative values and discovered that these two parameters could be used to characterize three rice cultivars with different RSA phenotypes (Fig. 6). These findings suggest that *θ*_*d*_ and log σ_*w*_ are useful parameters to compare the RSA of different varieties.

RGA is one of the most popular parameters in the vertical RSA phenotype (Oyanagi et al. 1993; Kato et al. 2006; Kitomi et al. 2015, 2020; An et al. 2017; Alahmad et al. 2019). The basket method of measuring the RGAs of individual roots is the most informative approach (Oyanagi et al. 1993; Kato et al. 2006). However, the basket method estimates RGA under the assumption that the roots are straight. In this study, we have shown that RGA calculated under this assumption is affected by distance in RGA measurement (distance between the planting site and point of measurement of RGA) and the horizontal parameter log σ_*w*_ (Fig. 8). For example, when *θ*_*d*_ is 60° and the distance in RGA measurement is 10 cm, RGA is overestimated by approximately 12% (7.1/60) in log σ_*w*_ of 3 and underestimated by 13% (7.8/60) in log σ_*w*_ of 2 (Fig. 8b, Online Resource 1). The difference is 25%, and this error will become problematic when making exact measurements of roots with various values of log σ_*w*_. In three rice varieties (IR64, Dro1-NIL, and KP), the log σ_*w*_ values of radicles and crown roots were close to 3 and 2, respectively (Fig. 6b). These findings suggested that RGA calculated under that assumption that the roots are straight would be overestimated in radicles and underestimated in crown roots.

To our knowledge, few studies have calculated the horizontal RSA phenotype while focusing on the shapes of individual roots. We assume that this is because most root phenotypings have been performed in 2D (Atkinson et al. 2019; Jia et al. 2019; Wang et al. 2019; Martins et al. 2020) and even if it has been conducted in 3D, root distribution parameters such as root length density and horizontal percentage area coverage have been focused on, whereas individual root shapes have not (Chen et al. 2019; Gao et al. 2019a, b; Teramoto and Uga 2020). In this study, we calculated log σ_*w*_ as a horizontal parameter and showed that three rice varieties displayed varietal differences of log σ_*w*_. These findings also suggest that log σ_*w*_ may be genetically alterable. However, we found correlation between *θ*_*d*_ and log σ_*w*_ in the three rice varieties (Fig. 6c), indicating simple links of vertical and horizontal phenotypes. Further study is required to ascertain whether *θ*_*d*_ and log σ_*w*_ are independent.

We have considered that our methodology will be used in studying the biological process of root development. In this study, we found that *θ*_*d*_ and log σ_*w*_ differed between radicles and crown roots (Figs. 6a and b), suggesting that radicles and crown roots may have different mechanisms of physiological development. Many studies have carried out root physiological experiments, such as gravitropism tests and stress response tests, on radicles (Uga et al. 2013; Kitomi et al. 2015; Sun et al. 2018). This is because the radicle is the first root to develop and it is easy to control both the angle and elongation of developing roots. Our methodology enables the physiological differences between seed and crown roots to be quantified, and this is expected to bring new insights.

The methodology of this study has three limitations: (1) it is not designed for roots that grow upward. We observed few roots extending from the lower soil layer to the upper soil layer. We considered that such roots may appear if the variety, plant species, or pot size are changed. If the conditions change, the equation should be reexamined. (2) It cannot accommodate roots that change their shape during cultivation; for example, when a plant is grown under normal conditions during the first half of its growth and then moved to grow under stress conditions. In these cases, the equation should be changed by introducing a variable representing time (Pagès et al. 2004). (3) As our model ignores lateral roots, it cannot be used to either evaluate or simulate whole root density in the soil. In other words, a simulation study of water and nutrient uptake (Postma et al. 2017; Daly et al. 2018; Takahashi and Pradal 2021) could not be conducted. Conversely, root distribution in the soil could be simulated, as shown in the case of RSA breeding by modifying the shapes of radicles and crown roots (Uga et al. 2013; Kitomi et al. 2020).

We named this script for calculating *θ*_*d*_, *θ*_*p*_, and log σ_*w*_ “RSAparam3D”. RSAparam3D are available from the GitHub repository (https://github.com/st707311g/RSAparam3D.git). It was developed as an extension module of RSAtrace3D (Teramoto et al. 2021). Thus, it is very easy to calculate parameters; RSAtrace3D automatically calculates *θ*_*d*_, *θ*_*p*_. and log σ_*w*_ when the root system is vectorized.

## Conclusion

We have demonstrated RSAparam3D, RSA parameterization software that calculates vertical and horizontal RSA phenotypes separately. With root system data vectorized by RSAtrace3D, various parameters can be calculated automatically. As RSAparam3D can be applied not only to phenotypic analysis but also to simulation analysis, it is expected to further promote root research.

## Supporting information

Online Resource 1

## Acknowledgments

This work was supported by JST CREST Grant Number JPMJCR17O1, Japan.

## Declarations

### Funding

This work was supported by JST CREST Grant Number JPMJCR17O1, Japan.

### Competing interests

The authors declare that they have no conflicts of interest.

### Availability of data and materials

The plant materials and raw data of the figures used in this study are available from the corresponding author on reasonable request.

### Authors’ contributions

Shota Teramoto designed the study, acquired the data, constructed and evaluated the code for calculating RSA parameters, and wrote the manuscript. Yusaku Uga coordinated the project and revised the manuscript.

### Code availability

Available from the GitHub repository (https://github.com/st707311g/RSAparam3D.git) or the project homepage (https://rootomics.dna.affrc.go.jp/en/).

### Ethics approval

Not applicable.

### Consent to participate

Not applicable.

### Consent for publication

Not applicable.

## Notes

### Competing Interest Statement

The authors have declared no competing interest.

